# Label-free 4D holotomography with depth-adaptive segmentation for quantitative analysis of lipid droplet dynamics in hepatic organoids

**DOI:** 10.64898/2026.04.01.707237

**Authors:** Jimin Cho, Hoyeon Lee, ChulMin Oh, Juyeon Park, Sujin Park, Bon-Kyoung Koo, YongKeun Park

## Abstract

**Significance:** Quantifying lipid droplet (LD) remodeling in 3D hepatic organoids is often limited to endpoint staining or phototoxic live fluorescence imaging, thereby obscuring droplet-level kinetics.

**Aim:** We aimed to develop a label-free method to track LD dynamics in living hepatic organoids under different fatty-acid loads.

**Approach:** Time-lapse 3D refractive-index tomograms were acquired using holotomography and analyzed with a depth-adaptive, multi-threshold segmentation pipeline to quantify LD number, volume, sphericity, and refractive-index–derived concentration and dry mass at single-droplet resolution.

**Results:** Oleic acid and linoleic acid induced LD accumulation while preserving organoid integrity, whereas palmitic acid triggered rapid structural collapse. Despite increases in total LD burden under both oleic acid and linoleic acid, droplet-level dynamics diverged: oleic acid produced volume-dominated accumulation via enlargement of fewer LDs and increased size heterogeneity, whereas linoleic acid produced number-dominated accumulation via sustained increases in LD number, yielding a more uniform population of small droplets.

**Conclusions:** Label-free holotomography with depth-adaptive analysis enables non-invasive, longitudinal, and multi-scale quantification of LD dynamics in intact organoids and reveals fatty-acid– dependent temporal modes of lipid storage.

**Statement of Discovery:** We developed a label-free, longitudinal 3D holotomography framework with depth-adaptive lipid droplet segmentation that quantifies single-droplet dynamics in living mouse hepatic organoids. Using this platform, we found that oleic acid and linoleic acid induce LD accumulation via distinct strategies—oleic acid via droplet enlargement and linoleic acid via sustained increases in droplet number—while palmitic acid rapidly compromises organoid integrity.

## 1 Introduction

Lipid droplets (LDs) are dynamic intracellular organelles that play central roles in lipid storage, energy homeostasis, and cellular stress responses^1^. Beyond serving as passive lipid reservoirs, LDs actively participate in metabolic regulation, oxidative stress buffering, and organelle crosstalk, and their dysregulation is implicated in a broad spectrum of diseases, including metabolic dysfunction-associated fatty liver disease (MAFLD), alcoholic liver disease (ALD), diabetes, cancer, and neurodegeneration. In hepatic systems, both the amount and the organization of LDs—such as their size, number, and spatial distribution—are closely linked to disease phenotype and progression^2,3^.

Recent advances in three-dimensional (3D) liver organoid models have provided powerful platforms for studying hepatic metabolism in a physiologically relevant context^4–8^. Compared with two-dimensional cell cultures, organoids recapitulate tissue architecture, cell–cell interactions, and metabolic heterogeneity, enabling more faithful modeling of lipid accumulation and steatosis. However, quantitative analysis of LD dynamics in organoids remains challenging. LD accumulation in organoids is inherently heterogeneous and highly dynamic, evolving over hours to days through coordinated processes of droplet biogenesis, growth, remodeling, and redistribution. Capturing these processes requires longitudinal, volumetric imaging with sufficient resolution and minimal perturbation.

Conventional LD assays rely primarily on fluorescence microscopy using lipid dyes or antibody labeling^9^. While widely used, these approaches suffer from several limitations when applied to 3D organoids [Fig. 1(a)]. Fixation and staining disrupt native lipid metabolism and preclude longitudinal measurements^10,11^. Live-cell fluorescence imaging is constrained by phototoxicity, photobleaching, and limited penetration depth, often restricting analysis to 2D projection images or endpoint measurements^4,12^. As a result, most existing studies characterize LD accumulation using static, population-averaged metrics that obscure droplet-level heterogeneity and temporal organization^4,13^.

**Fig. 1.**
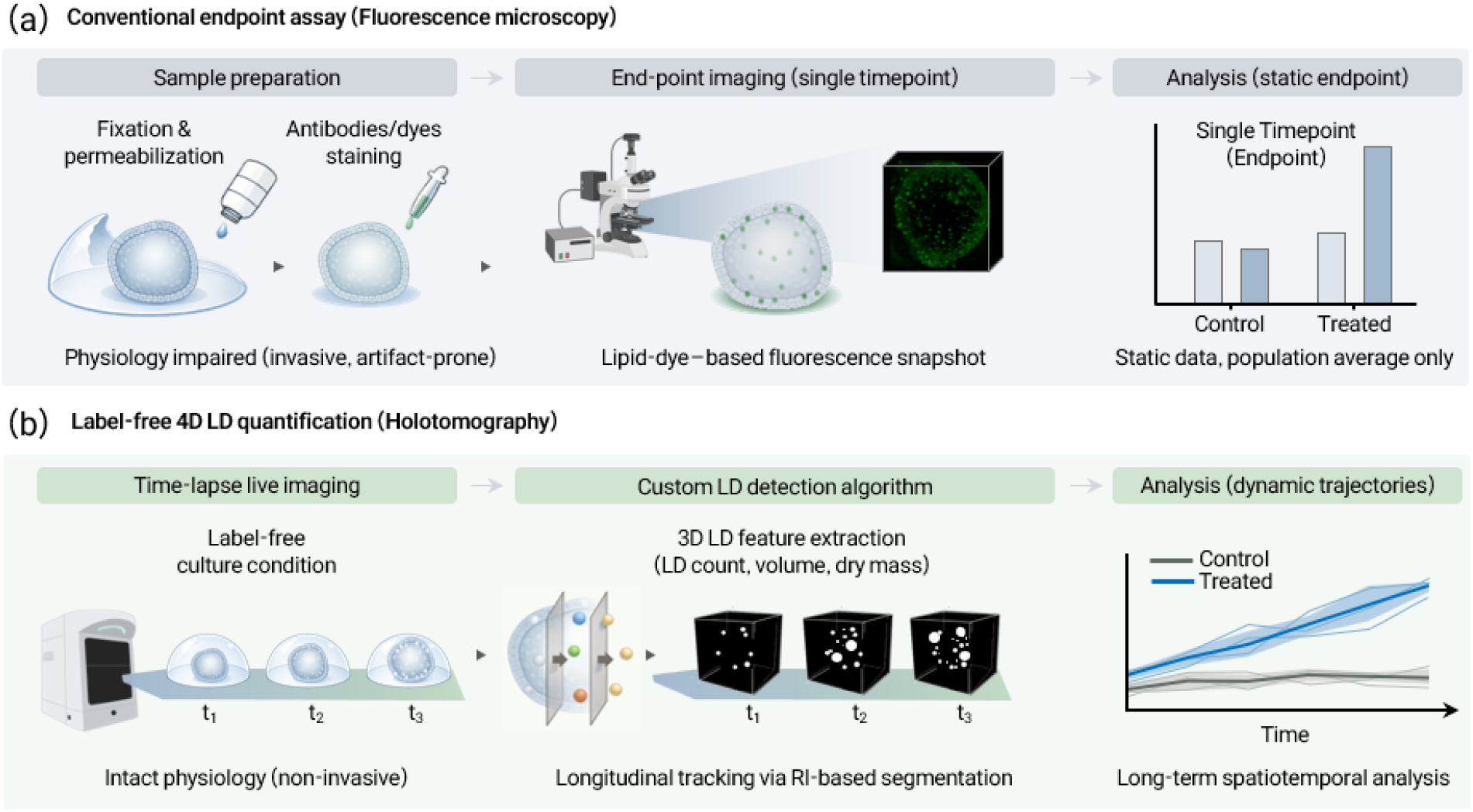
Conceptual comparison between conventional fluorescence-based LD analysis and the proposed label-free 4D HT approach. (a) Conventional fluorescence-based assays rely on fixation, permeabilization, and staining with dyes or antibodies, which perturb cellular physiology and typically limit analysis to static endpoint measurements. (b) The proposed workflow enables non-invasive, time-lapse 3D imaging of live organoids while preserving intact physiology, allowing longitudinal, single-droplet–resolved quantification and dynamic trajectory analysis.

Label-free optical imaging methods based on refractive index (RI) contrast provide an alternative. Holotomography (HT), a 3D quantitative phase imaging technique, reconstructs quantitative 3D RI distributions from multiple angle-resolved intensity measurements, enabling non-invasive imaging of intact, living samples^14–25^. Because neutral lipids exhibit intrinsically high RI relative to the surrounding cytoplasm, HT has been used to visualize LDs in single cells without exogenous labels. Extending this capability to thick, multicellular organoids, however, introduces new challenges. Depth-dependent RI variations, optical aberrations, and organoid structural heterogeneity complicate reliable LD identification, rendering simple global-threshold segmentation strategies insufficient. Moreover, most existing LD analyses—whether fluorescence-based or label-free—focus on organoid-or cell-level bulk metrics, such as total lipid content or average droplet size. Such measurements cannot disentangle whether lipid accumulation arises from increases in droplet number, droplet size, or changes in spatial organization. As a result, distinct lipid-storage strategies driven by different fatty acids may appear similar at the population level, despite fundamentally different underlying dynamics.

Here, we present a quantitative, label-free framework for tracking LD dynamics in living mouse hepatic organoids (mHOs) using HT combined with depth-adaptive segmentation and longitudinal 3D quantification [Fig. 1(b)]. By resolving LDs across depth layers and imaging the same organoid over time, our approach enables analysis of LD number, size, and RI-derived biophysical properties at single-droplet resolution within intact organoids. Using this framework, we systematically compare LD accumulation dynamics induced by unsaturated free fatty acids (FFAs)— oleic acid (OA) and linoleic acid (LA)—and contrast them with the lipotoxic response elicited by saturated palmitic acid (PA). We show that OA- and LA-induced lipid accumulation proceeds through distinct droplet-level strategies.

## 2. Method

### 2.1 Organoid preparation

MHOs were obtained from a commercial source (STEMCELL Technologies) and maintained in HepatiCult™ Organoid Growth Medium (Mouse; STEMCELL Technologies) according to the manufacturer’s protocol. Organoids were embedded in Matrigel domes and cultured under standard conditions (37 °C, 5% CO_2_). For passaging, organoids were released from Matrigel by gentle mechanical dissociation, collected by centrifugation (300 × g, 5 min), and resuspended in fresh Matrigel for re-plating. Organoids were routinely subcultured at a split ratio of 1:3 to 1:6 every 5–7 days, depending on growth rate and density.

### 2.2 Live imaging of hepatic organoids

To enable non-invasive visualization of LDs in living mHOs, we employed the HT system (HT-X1, Tomocube Inc.) that reconstructs 3D RI distributions from multiple intensity measurements acquired under spatially programmed illumination patterns [Fig. 2(a)]. Raw intensity images acquired under multiple illumination angles using a digital micromirror device (DMD) are processed with an HT reconstruction algorithm to generate 3D RI tomograms^26–28^. Organoids were imaged in 96-well plates under physiological conditions (37°C, 5% CO_2_) using an on-stage incubation system. Time-lapse imaging was conducted at 6-h intervals for up to 24 or 42 h, depending on the experimental condition. By exploiting intrinsic optical contrast rather than exogenous labels, HT allows volumetric imaging of intact organoids while preserving native cellular physiology^29,30^.

**Fig. 2.**
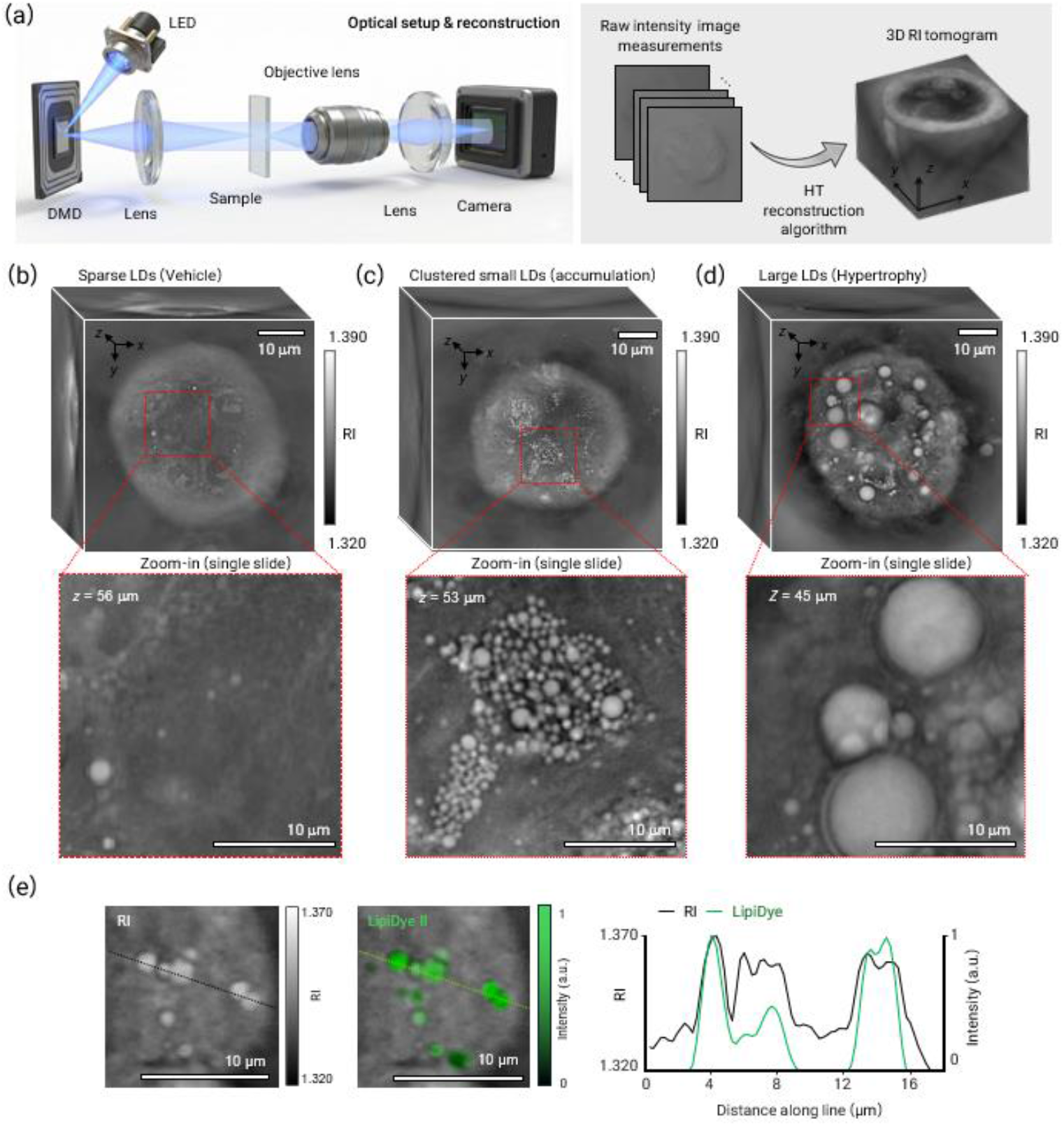
Label-free HT imaging and representative visualization of LD accumulation in mHOs. (a) Schematic of the HT optical setup and reconstruction pipeline. (b)–(d) Representative label-free 3D RI tomograms of live mHOs acquired by HT, illustrating distinct LD accumulation phenotypes: (b) an organoid with sparse LDs (vehicle control); (c) an organoid exhibiting accumulation of numerous small LDs; and (d) an organoid dominated by enlarged LDs. Enlarged insets show internal cross-sectional views at the indicated axial depths, highlighting LD-rich regions within the organoids. (e) Correlative fluorescence validation demonstrating spatial colocalization between high-RI structures detected by HT and LDs stained with LipiDye II. Line profiles across representative LDs show concordant RI and fluorescence signals, supporting the specificity of RI contrast for LD identification.

Reconstructed RI tomograms revealed heterogeneous LD accumulation patterns within mHOs, which could be readily visualized through optical sectioning of the 3D datasets [Figs. 2(b)–(d)]. It is known that the RI values of LDs (*n* > 1.37) are distinctively higher than that of cytoplasm^9,31,32^. Representative examples included organoids containing only sparse LDs under vehicle conditions [Fig. 2(b)], organoids exhibiting numerous small LDs distributed throughout the cytoplasm [Fig. 2(c)], and organoids dominated by a smaller number of enlarged LDs occupying substantial intracellular volume [Fig. 2(d)] under FFA treatment. Enlarged cross-sectional views at different axial depths further highlighted the 3D morphology and spatial distribution of LD-rich regions within intact organoids.

To verify that high-RI structures detected by HT correspond to LDs, we performed correlative fluorescence imaging using LipiDye II. High-RI inclusions observed in HT tomograms spatially colocalized with fluorescently labeled LDs [Fig. 2(e)]. Line-profile analysis across representative droplets showed concordant RI and fluorescence intensity peaks, supporting the specificity of RI contrast for LD identification. Together, these results establish HT as a robust platform for label-free, 3D visualization of LD morphology and distribution in live hepatic organoids, providing the basis for subsequent quantitative analyses.

### 2.3 Organoid segmentation algorithm

Organoid segmentation was adapted from a previously reported MP-HT–based pipeline^33^ with minor modifications. Briefly, 3D RI volumes were first down-sampled to reduce high-frequency noise, denoised using median filtering, and subsequently up-sampled to the original resolution. An initial organoid mask was then obtained by adaptive Otsu thresholding followed by selection of the largest connected component. To refine the organoid boundary, a slice-wise, two-pass adaptive morphological closing operation was applied within a cropped region of interest. For each axial slice, the minimum structuring-element radius that achieved at least a 10% filling gain was selected, allowing robust closure of boundary gaps while minimizing over-segmentation. The resulting mask was subsequently thinned and resampled to match the original RI voxel dimensions. The lumen mask was derived by subtracting the refined organoid mask from its hole-filled counterpart, followed by retention of the largest connected component. Organoid and lumen volumes were computed independently at each time point. The final organoid volume was defined as the epithelial compartment, calculated as the total organoid volume excluding the lumen volume.

### 2.4 LD segmentation algorithm

While HT enables label-free 3D visualization of LDs in mHOs, reliable quantitative analysis requires accurate extraction of LDs from reconstructed RI tomograms. In contrast to single-cell imaging, LD segmentation in thick organoids is challenged by depth-dependent RI variations, heterogeneous background structures, and reduced LD-to-background contrast, rendering global threshold–based approaches insufficient (Fig. S1 in the Supplementary Material). A custom rule-based segmentation algorithm was implemented in MATLAB (MathWorks) to extract LDs from 3D RI tomograms, as summarized schematically in Fig. 3. The algorithm was designed to handle depth-dependent contrast variations via multiple RI-contrast thresholds and to suppress heterogeneous background structures via morphological filtering.

**Fig. 3.**
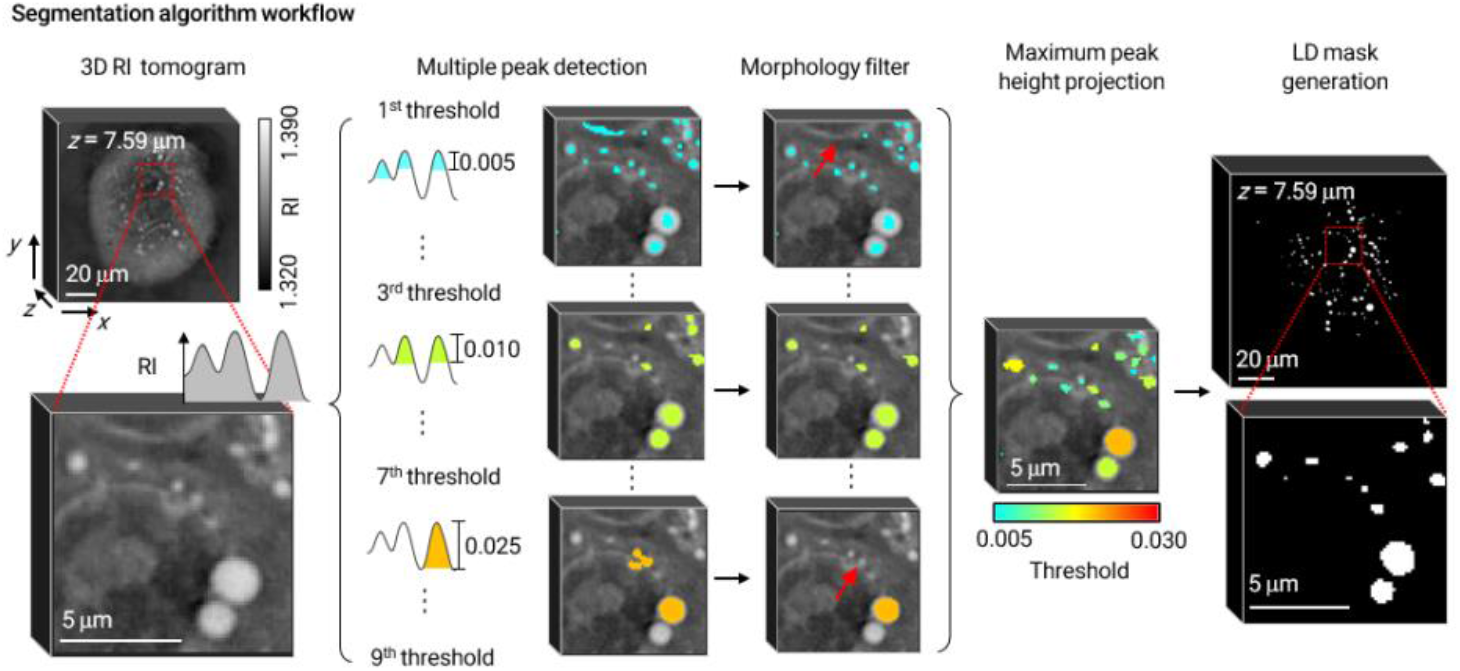
Schematic overview of the LD segmentation algorithm workflow. Candidate LDs are identified across multiple RI contrast thresholds (Δ*n* = 0.005–0.03) to account for heterogeneity in LD intensity and depth-dependent contrast variations. Detected objects are subsequently refined using morphological filtering criteria to exclude irregular or non-spherical structures, yielding 3D binary LD masks.

LDs were segmented within an organoid-bounded cropped volume (bounding box from the organoid mask) to reduce background interference and peripheral truncation. LD candidates were detected as 3D regional maxima across nine RI contrast thresholds (Δ*n* = 0.005, 0.007, 0.010, 0.012, 0.015, 0.017, 0.020, 0.025, 0.030), and filtered using 3D morphological criteria (compactness < 4.4, solidity > 0.5, aspect ratio < 2.0) followed by a 2D Feret diameter ratio constraint (max/min < 1.5) to retain near-spherical objects. Candidates from all thresholds were merged and 3D hole filling was applied to generate an initial LD mask. To reduce false positives while preserving large droplets, a size-adaptive refinement was performed using 2D MIP (Maximum Intensity Projection) major-axis length: objects with major axis ≤ 5 pixels were restricted to voxels detected at Δ*n* ≥ 0.010, whereas larger objects were retained without truncation. The resulting binary volume was used as the final LD mask for downstream analyses.

### 2.5 Algorithm validation

Application of the algorithm to reconstructed RI datasets demonstrated consistent LD detection across axial depths within intact organoids [Fig. 4(a)]. Layer-resolved comparisons of raw RI slices and the corresponding LD masks showed that LD-like structures could be extracted throughout the organoid volume, including regions where local background RI varied substantially. These results indicate that the adaptive thresholding strategy effectively mitigates depth-related contrast heterogeneity inherent to organoid imaging.

**Fig. 4.**
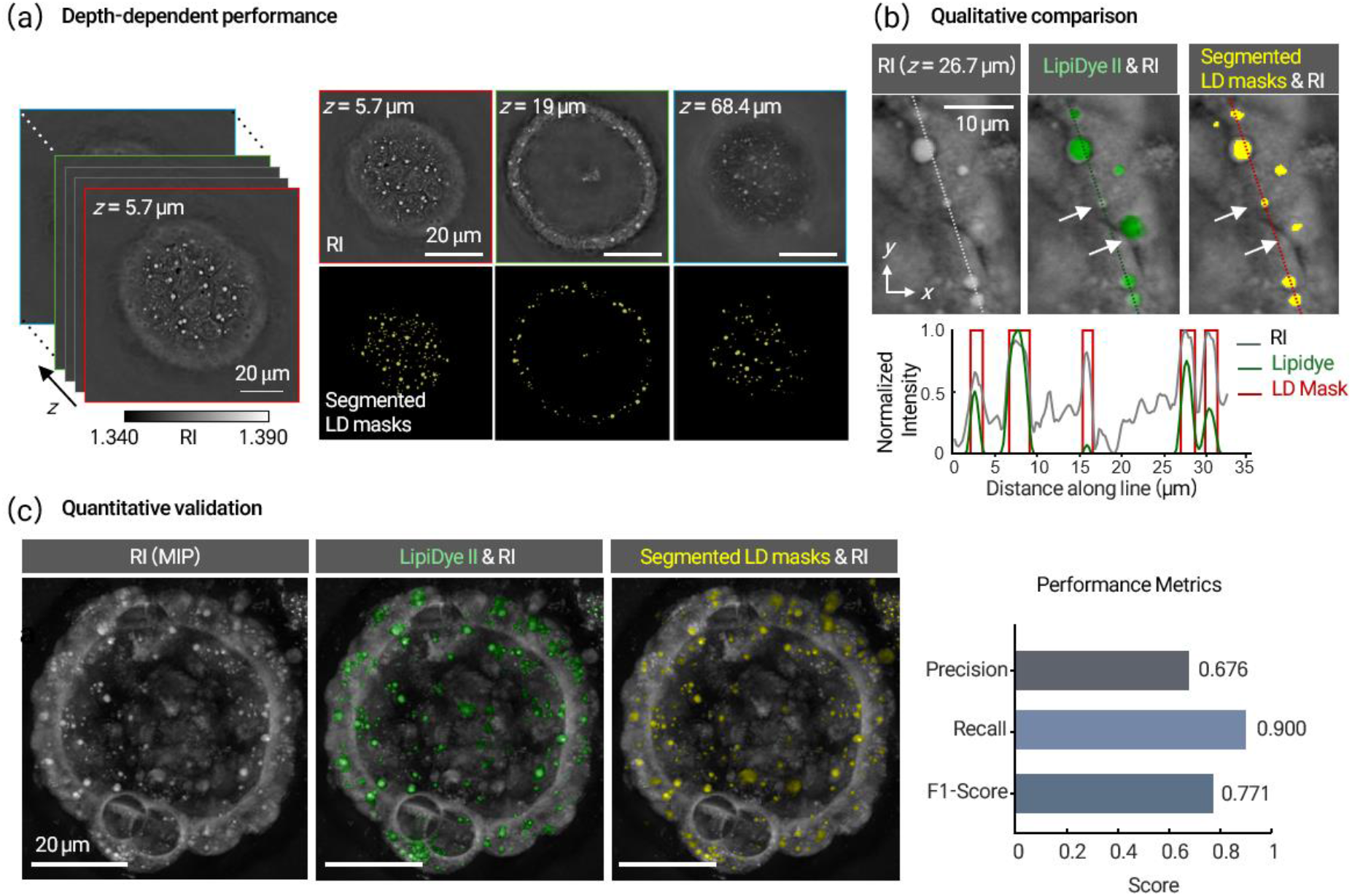
Validation of a depth-adaptive algorithm for 3D LD segmentation. (a) Representative layer-resolved segmentation results at different axial depths within a hepatic organoid, showing reconstructed RI slices and corresponding LD masks, illustrating robust LD detection across depth. (b) Representative comparison between RI images, fluorescence images acquired using LipiDye II, and the corresponding RI-based LD segmentation outputs. Line profiles along the indicated line show correspondence among RI, LipiDye, and segmented masks. (c) Quantitative validation of RI-based LD segmentation using correlative fluorescence data displayed as MIPs.

To qualitatively compare RI-based LD segmentation with conventional fluorescence (FL) imaging, we performed correlative imaging using the neutral lipid dye LipiDye II (Funakoshi) [Fig. 4(b)]. LipiDye II was diluted in organoid culture medium to a final concentration of 1 µM, and organoids were incubated with the staining solution for 1 h at 37°C in a humidified 5% CO_2_ atmosphere prior to imaging. FL images exhibited depth-dependent signal attenuation and blurred droplet boundaries, particularly at increased imaging depths, whereas RI-based segmentation provided label-free delineation of LD-like structures across the full 3D volume. Line-profile analysis further showed spatial correspondence between RI peaks, fluorescence signals, and segmented LD masks, supporting the specificity of RI contrast for LD detection.

Quantitative validation of the segmentation results was performed using correlative FL data [Fig. 4(c)]. FL image volumes were linearly rescaled to a fixed intensity range (30–300) and then binarized to generate reference FL masks. Connected-component analysis was applied to the binarized FL volumes to identify individual fluorescent objects. Because of the limited axial resolution of confocal fluorescence imaging in thick organoids, the small physical size of many LDs (often spanning only a few voxels), and residual spatial misregistration between HT and FL datasets, accurate instance-level correspondence between individual LDs could not be reliably established. Consequently, segmentation accuracy was assessed on a volume-weighted basis rather than at the single-object level. To accommodate these constraints, a permissive overlap criterion was adopted: an HT-derived LD label and an FL-derived label were considered a true positive (TP) if they shared at least one voxel. Labels without overlap were classified as false positives (FP) or false negatives (FN) accordingly. This criterion was chosen to minimize penalization arising from axial blurring and registration uncertainty in FL images, while still providing a conservative assessment of volumetric correspondence. Precision and recall were defined as follows: 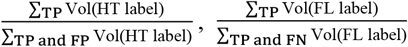. The F1-score was calculated as the harmonic mean of precision and recall: 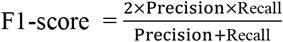. Segmentation performance was evaluated in 3D using label-volume-weighted metrics, yielding a precision of 0.676, recall of 0.900, and an F1-score of 0.771. These values reflect high sensitivity for LD detection while acknowledging residual false-positive detections arising from high-RI non-lipid structures within organoids.

These results establish a depth-adaptive, morphology-aware segmentation framework that enables robust extraction of LD-like structures from HT-derived RI tomograms in intact organoids. This capability forms the computational foundation for the longitudinal, single-organoid, and single-droplet analyses presented in the following sections.

### 2.6 Free fatty acid-induced LD accumulation

To investigate the diversity of LD accumulation dynamics underlying hepatic steatosis, we selected three FFAs—OA, LA, and PA—which differ in their degree of saturation and are known to induce distinct biophysical and metabolic responses in hepatocytes and liver organoid models^4,34–37^.

OA, LA, and PA powders were dissolved in ethanol (EtOH) to prepare 100 mM stock solutions. To generate FFA–BSA conjugates, FFA stocks were mixed with 10% (w/v) bovine serum albumin (BSA) solution at a 1:9 (v/v) ratio and incubated at room temperature with gentle agitation until fully conjugated. Vehicle controls were prepared by mixing an equivalent volume of EtOH with 10% BSA solution following the same procedure. FFA–BSA conjugates were diluted in organoid culture medium to final FFA concentrations of 200 µM or 400 µM immediately prior to use.

Following passaging, mHOs were allowed to stabilize for 5 h and were then exposed to individual FFAs. Beginning 30 min after FFA addition, organoids were imaged by HT at 6-h intervals to capture LD dynamics over time (Fig. 5). The resulting 3D RI datasets were processed using the depth-adaptive LD segmentation algorithm described above, yielding 3D LD masks for each time point. This pipeline enabled longitudinal, single-organoid and single-droplet–resolved quantification of LD number, size, and RI-derived properties within intact, living organoids.

**Fig. 5.**
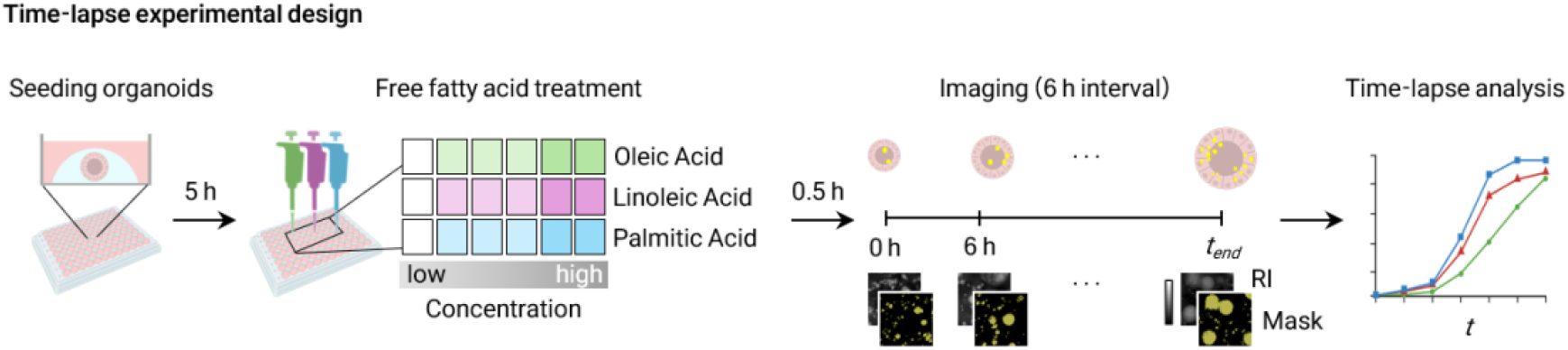
Experimental workflow of free fatty acid (FFA)–induced LD accumulation in hepatic organoids. Schematic overview of the experimental and analytical workflow for longitudinal, label-free quantification of LD accumulation in mHOs following FFA treatment. Organoids were imaged by HT at 6-h intervals and analyzed using automated LD segmentation and time-resolved quantitative metrics.

### 2.7 Quantitative feature extraction

All quantitative features were computed using the native voxel dimensions of each image and extracted in MATLAB using the regionprops3 function. Organoids exhibiting loss of lumen integrity, necrotic core formation, or severe structural collapse were classified as low-viability and excluded from all quantitative analyses (Fig. S2 in the Supplementary Material). In addition, particularly in LA-treated groups, organoids showing excessive axial thickening or lateral expansion beyond the field of view were excluded to prevent volumetric truncation artifacts and ensure accurate holotomographic reconstruction. For all quantitative analyses, *n* denotes the number of individual organoids. Sample sizes were as follows: vehicle, *n* = 4 organoids from 3 wells; OA 200 µM, *n* = 9 organoids from 3 wells; OA 400 µM, *n* = 11 organoids from 5 wells; LA 200 µM, *n* = 3 organoids from 3 wells; and LA 400 µM, *n* = 3 organoids from 3 wells.

LD volume was calculated by multiplying the number of voxels in the segmented LD mask by the voxel size in the *x, y*, and *z* dimensions. LD surface area was estimated from the 3D binary LD masks using a voxel-based surface area metric implemented in MATLAB. For each segmented LD, the effective lipid concentration *c*_*LD*_ was computed as 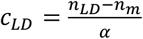, where *n*_*LD*_ is the mean RI of the LD, *n*_*m*_ is the RI of the surrounding medium, and *α* is the RI increment of lipids^38^. We adopted *α* = 0.135 mL/g, which is widely used for neutral lipids in HT-based quantification of LDs, including live hepatocyte studies and fatty-acid perturbation experiments^39^. This value reflects the RI increment of neutral lipid cores (primarily triacylglycerols and cholesteryl esters) and has been consistently applied in prior work to estimate lipid concentration and dry mass from RI contrast^38,40^. The dry mass of each LD was then calculated as *M*_*LD*_ = *c*_*LD*_ × *V*_*LD*_, where *V*_*LD*_ is the LD volume obtained from 3D segmentation. LD sphericity (Ψ) was calculated to quantify droplet shape using the standard definition: 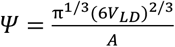, where A denotes LD surface area. A sphericity value of 1 corresponds to a perfect sphere, with lower values indicating increasing deviation from spherical morphology.

## 3. Results

### 3.1 Longitudinal, label-free tracking of LD accumulation in hepatic organoids

Figure 6(a) presents representative MIP images of mHOs under different FFA conditions, acquired every 6 h over a 24-h period. These time-lapse images provide a qualitative overview of how distinct FFA environments shape LD accumulation and organoid-level morphology. In vehicle-treated organoids, only sparse, small LDs are observed throughout the imaging period, with minimal temporal change, consistent with basal lipid turnover in the absence of exogenous FFAs^41,42^. OA-treated organoids exhibit progressive LD accumulation characterized by the emergence and growth of prominent LDs over time, while largely preserving overall organoid morphology. In contrast, LA-treated organoids show a rapid and widespread increase in the number of small LDs distributed throughout the organoid volume. PA-treated organoids display a markedly different behavior. At early time points, LDs are detectable; however, progression over time is accompanied by pronounced morphological alterations, including surface irregularities, membrane blebbing [red arrow in Fig. 6(a)], and loss of compact organoid architecture. These features indicate progressive structural destabilization under saturated fatty-acid loading and contrast sharply with the relatively preserved morphology observed under unsaturated FFA (OA, LA) treatment.

**Fig. 6.**
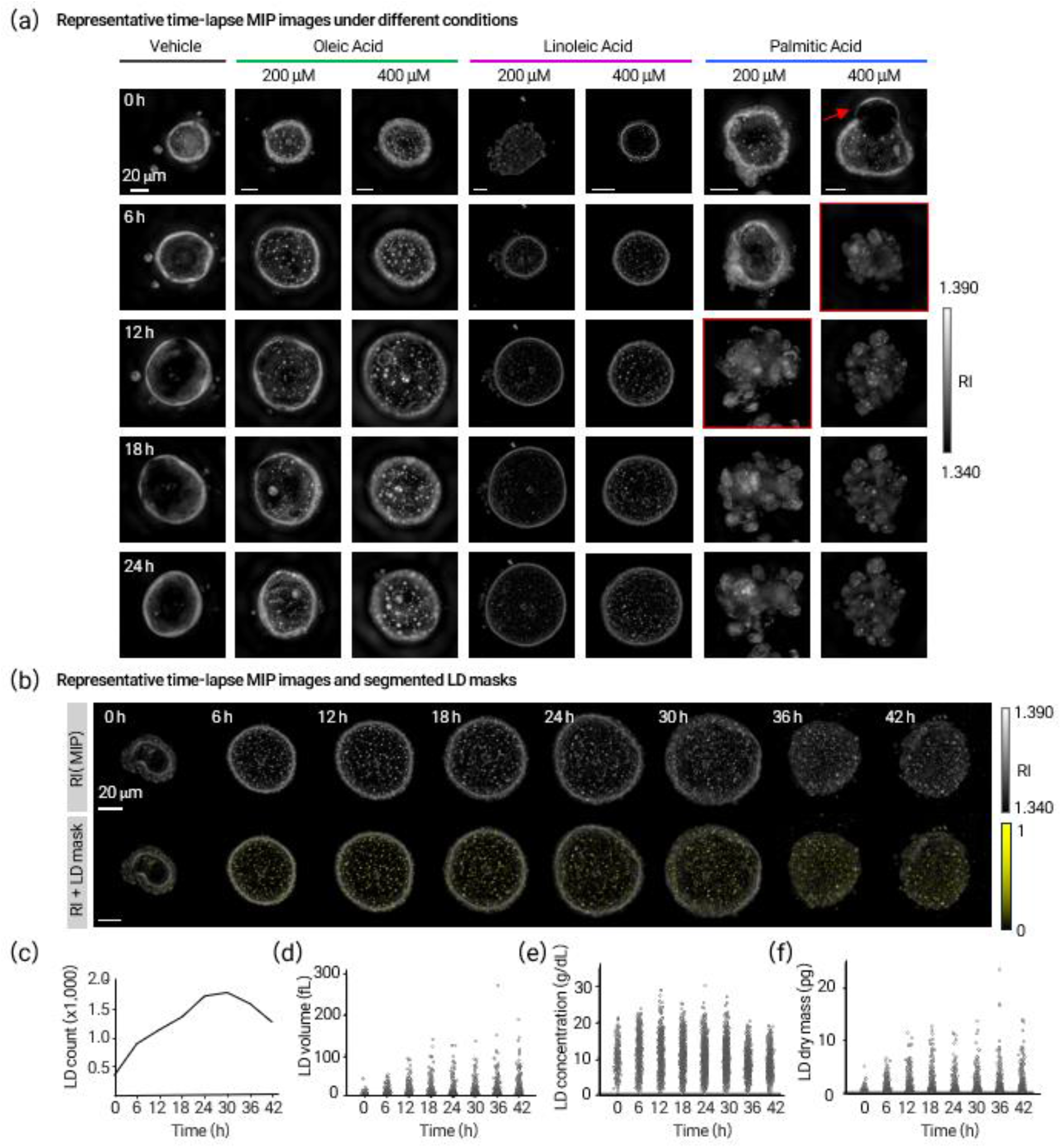
Representative time-lapse analysis of FFA–induced LD accumulation in hepatic organoids. (a) Representative time-lapse MIP images of mHOs treated with vehicle control, Oleic Acid (OA; 200 and 400 μM), Linoleic Acid (LA; 200 and 400 μM), and Palmitic Acid (PA; 200 and 400 μM). Organoids were imaged every 6 h for 24 h starting 30 min after FFA addition. Red boxes highlight time points exhibiting pronounced organoid disintegration, and arrows indicate membrane blebbing observed under PA treatment. (b) Representative time-lapse MIP images with segmented LD masks overlaid (yellow) of a live mHO treated with 200 μM LA, acquired every 6 h for up to 42 h after FFA exposure. (c)–(f) Single-droplet quantitative analysis corresponding to the organoid shown in (b), showing the temporal evolution of the (c) LD count (number of LDs); (d) LD volume (fL); (e) LD concentration (g/dL); and (f) LD dry mass (pg). In panels (d)–(f), each dot corresponds to an individual LD at a given time point.

Overlay of segmented LD masks on the time-lapse RI MIP images demonstrates that the proposed LD segmentation algorithm can be robustly applied across time points and treatment conditions for a representative LA– treated organoid [Fig. 6(b)]. LD masks are consistently detected throughout the full organoid volume rather than being confined to two-dimensional projections, indicating that the segmentation algorithm remains effective despite depth-dependent RI variations and changing surrounding structure during long-term live imaging.

### 3.2 Extraction of quantitative imaging parameters at the individual LD level

Building on the robust performance of the LD segmentation algorithm in time-lapse organoid datasets [Fig. 6(b)], we next evaluated whether the segmented LD masks enable reliable extraction of quantitative imaging parameters at the level of individual LDs. Using a representative LA–treated organoid as an example, multiple LD-level metrics were computed longitudinally from the segmented masks [Fig. 6(c)–(f)].

As shown in Fig. 6(c), the number of segmented LDs increased substantially over time, demonstrating that individual LDs can be consistently detected and enumerated across successive imaging time points. Specifically, the total LD count increased from 329 at 0 h to 1,227 at 42 h, reaching a maximum of 1,729 LDs at 30 h, indicating sustained LD generation over the course of the experiment. In parallel, the median LD volume increased from 2.0 fL at 0 h to 3.0 fL at 42 h [Fig. 6(d)], confirming that volumetric measurements derived from the segmented LD masks remain consistent and interpretable during long-term live imaging.

Beyond geometric descriptors, RI–derived parameters were also quantified at the individual LD level (see Sec. 2.7 for details of the feature extraction). Median LD concentration remained relatively stable over time, with a modest early increase followed by a partial decline and subsequent stabilization [Fig. 6(e)]. Importantly, LD dry mass was computed by combining LD volume with RI-derived LD concentration [Fig. 6(f)], demonstrating that higher-order, biophysically meaningful metrics can be derived directly from the segmented LD masks at single-droplet resolution. These profiles suggest that LD accumulation involves dynamic remodeling of both droplet count and volume, rather than uniform scaling of all droplets over time, reflecting their active role in lipid homeostasis and energy metabolism within organoids^43,44^.

Figures 6(c)–(f) establish that time-resolved LD segmentation enables systematic extraction of multiple quantitative imaging parameters—including LD number, size, and RI-derived properties—at the level of individual droplets in living organoids. This capability provides the technical foundation for subsequent population-level comparisons and fatty-acid–specific analyses of LD accumulation dynamics.

### 3.3 Population-level kinetics reveal fatty-acid–dependent LD accumulation in hepatic organoids

To determine how different FFAs modulate LD accumulation at the organoid scale, we quantified time-resolved LD and organoid-level metrics across all treatment conditions. For each mHO, total LD volume, total LD dry mass, organoid volume, and relative steatosis level were tracked longitudinally and expressed as fold changes relative to baseline (*t* = 0 h) [Figs. 7(a)–(d)].

**Fig. 7.**
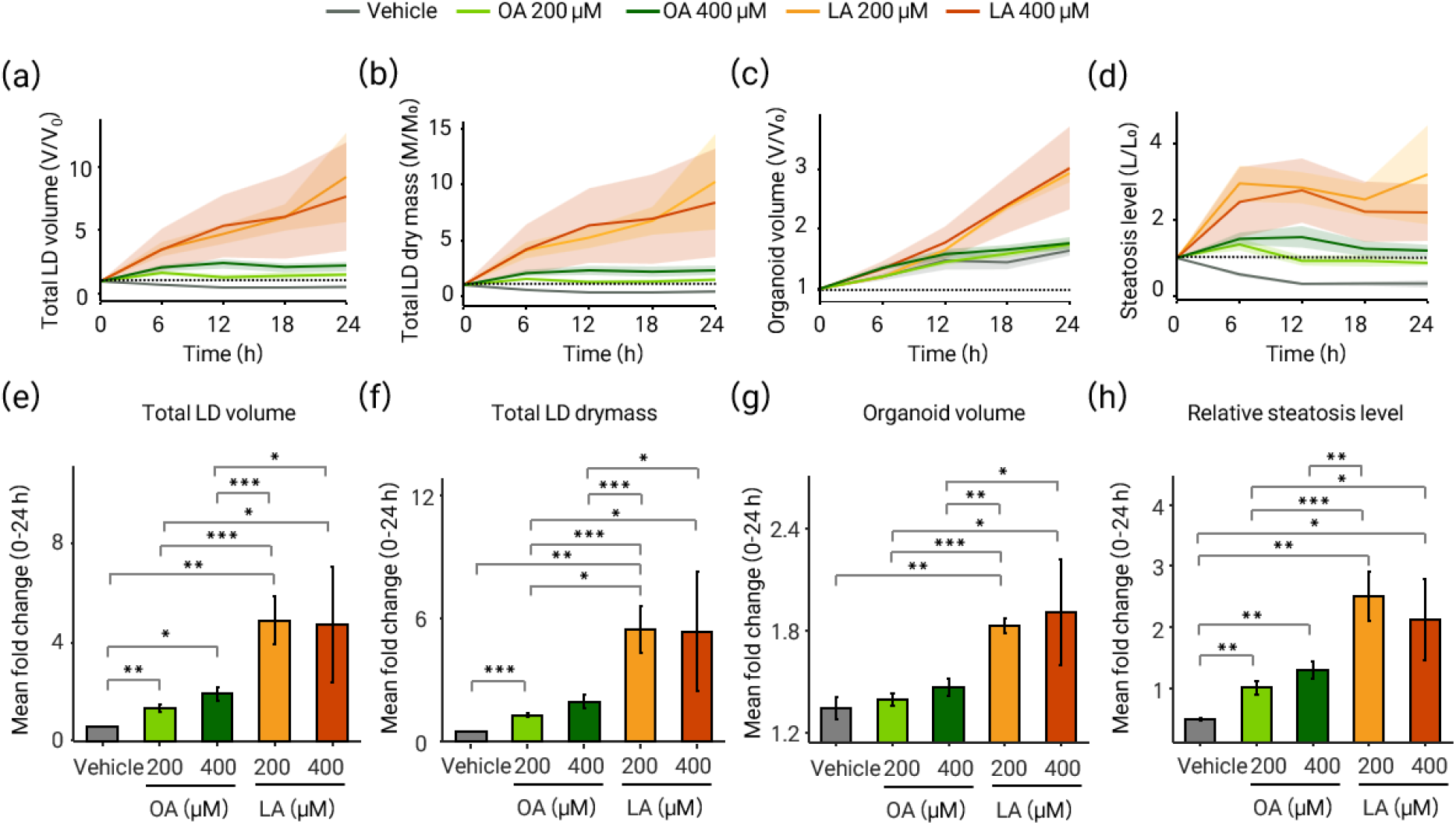
Label-free time-lapse analysis reveals distinct LD accumulation and responses of mHOs to FFAs. (a)–(d) Time-course profiles of (a) Total LD volume, (b) Total LD dry mass, (c) Organoid volume, and (d) Relative steatosis level, quantified per organoid from label-free HT data. All values are expressed as fold changes relative to the baseline at *t* = 0 h. Solid lines indicate group means, shaded regions represent ± s.e.m., and grey dashed lines denote baseline levels. (e)–(h) Bar plots summarizing per-organoid responses: for each organoid, fold-change values were averaged over 0–24 h to yield one time-averaged fold-change value, and bars show the group mean ± s.e.m. Overall group differences were assessed using one-way ANOVA. Pairwise differences between fatty-acid–treated groups are indicated (*\*p* < 0.05, \*\**p* < 0.01, \*\*\**p* < 0.001); comparisons without annotations were not statistically significant. Sample sizes: vehicle *n* = 4, OA 200 μM *n* = 9, OA 400 μM *n* = 11, LA 200 μM *n* = 3, LA 400 μM *n* = 3.

Time-course analysis showed minimal changes in all metrics in vehicle-treated organoids, consistent with basal lipid turnover under FFA-free control conditions^41,42,45^. In contrast, treatment with unsaturated fatty acids—OA and LA—induced pronounced increases in total LD volume and total LD dry mass over the 24-h observation period [Figs. 7(a), (b)]. These results are consistent with efficient incorporation of exogenous FFAs into neutral lipid stores and LD biogenesis^35,36,46,47^. Notably, LA-treated organoids displayed a sustained rise in total LD content across the full observation period, resulting in higher overall accumulation than OA at equivalent concentrations. This behavior is consistent with previous studies comparing OA- and LA-induced LD accumulation in two-dimensional hepatocyte cultures^48^. In contrast, OA-treated organoids reached an apparent plateau early in the time course, consistent with reports that OA-induced LD formation can occur within minutes of exposure^46^. Furthermore, organoid volume increased in all FFA-treated groups, with a markedly stronger effect under LA treatment [Fig. 7(c)]. This enlargement may reflect an injury-like swelling response, consistent with LA’s susceptibility to lipid peroxidation^49–51^ and the lumen collapse and necrotic core formation observed in a subset of LA-treated organoids (Supplementary Fig. S2). Relative steatosis level—defined as the ratio of total LD volume to organoid volume^4^—initially increased across all fatty-acid conditions [Fig. 7(d)]. At later time points, trajectories diverged, reflecting the combined effects of LD accumulation and concurrent changes in organoid volume. In LA-treated organoids, relative steatosis tended to plateau after ~12 h, consistent with LD accumulation occurring alongside substantial organoid expansion.

To enable statistical comparison across conditions while accounting for repeated measurements, per-organoid responses were summarized as time-averaged fold changes over the 0–24 h interval [Figs. 7(e)–(h)]. Both OA- and LA-treated organoids exhibited significantly higher mean fold changes in total LD volume and total LD dry mass compared with vehicle controls [Figs. 7(e), (f)]. Organoid volume and relative steatosis level also increased across treatments, highlighting their utility as an integrated, organoid-scale measure of lipid accumulation. Statistical significance was assessed by one-way ANOVA followed by Holm-adjusted unpaired, two-sided Welch’s *t*-tests for planned pairwise comparisons.

### 3.4 Divergent organoid responses to unsaturated and saturated fatty acids

In contrast to the robust yet non-disruptive lipid accumulation induced by unsaturated FFAs, treatment with the saturated fatty acid PA elicited a markedly different response. PA-treated organoids exhibited rapid morphological destabilization, including surface roughening, membrane blebbing, and eventual organoid disintegration at higher concentrations [Fig. 6(a)], consistent with its known lipotoxic effects reported in hepatocyte and liver model systems^34– 36,52^.

Specifically, at 200 μM PA, organoids initially maintained overall structure during early imaging but began to exhibit surface roughening and fragmentation after approximately 12 h. At 400 μM PA, membrane blebbing was apparent within 30 min and complete dissociation occurred by 6 h, reflecting acute mechanical and metabolic stress responses^53,54^. Due to this rapid structural deterioration, PA-treated organoids were excluded from quantitative kinetic comparisons.

### 3.5 Single–LD–resolved analysis reveals distinct accumulation modes under OA and LA treatment

The population-level analyses demonstrate that unsaturated FFAs drive robust LD accumulation in hepatic organoids while largely preserving overall organoid morphology. Although OA and LA both increase total LD burden at the organoid level, the metrics in Fig. 7 do not resolve how lipid accumulation is partitioned among individual droplets. We therefore performed single–LD–resolved analyses that dissect fatty-acid–specific accumulation strategies at higher resolution.

To determine how these increases in total LD burden arise at the level of individual droplets, we performed single– LD–resolved temporal analyses focusing on OA- and LA-treated organoids (Fig. 8). High-magnification MIP images revealed qualitatively distinct LD accumulation behaviors under OA and LA treatment at matched concentrations (400 μM) [Figs. 8(a), (b)]. OA-treated organoids exhibited progressive growth of a subset of LDs, leading to the emergence of larger droplets over time, whereas LA-treated organoids showed a widespread increase in the number of small LDs distributed throughout the organoid volume.

**Fig. 8.**
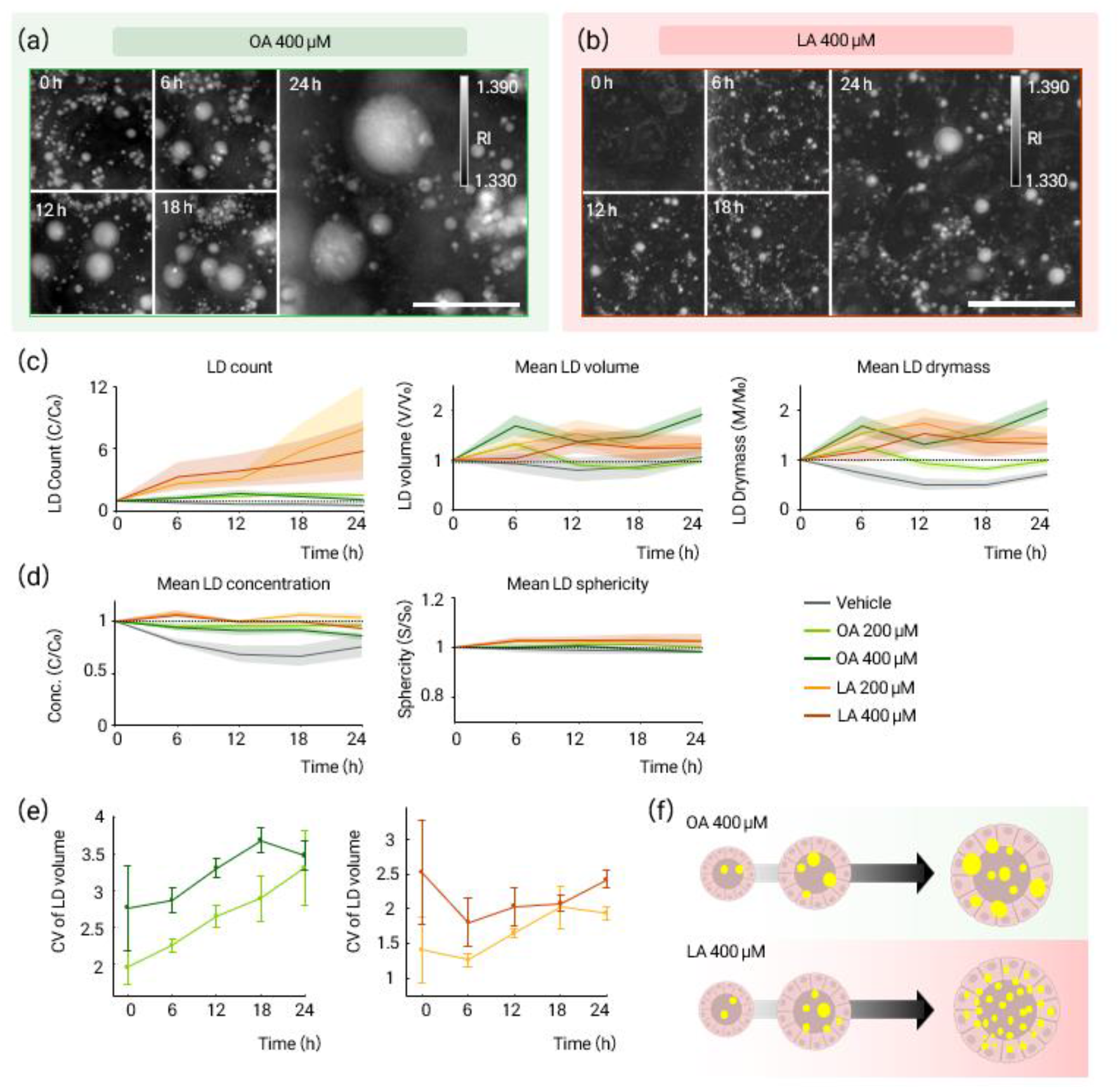
Single–LD–resolved temporal analysis reveals distinct lipid accumulation patterns under OA and LA treatment. (a)–(b) High-magnification MIP images of mHOs treated with (a) 400 μM OA or (b) 400 μM LA, showing time-dependent accumulation and remodeling of individual LDs. Images are shown at representative time points from 0 to 24 h after treatment (scale bars, 10 μm). (c)–(d) Time-course quantification of LD-level metrics across all experimental conditions (vehicle, OA 200 μM, OA 400 μM, LA 200 μM, and LA 400 μM): (c) LD count, mean LD volume, and mean LD dry mass; (d) mean LD concentration and mean sphericity. Metrics were computed per organoid and are expressed as fold changes relative to t = 0 h. Solid lines indicate group means and shaded regions indicate ± s.e.m.; dashed grey lines indicate baseline levels. (e) Temporal evolution of LD size heterogeneity, quantified as the CV of individual LD volumes. For each time point, CV values were calculated per organoid and subsequently averaged within each treatment group (mean ± s.e.m.). (f) Conceptual summary illustrating contrasting LD accumulation patterns observed under OA and LA treatment. OA exposure is associated with preferential enlargement of fewer LDs, whereas LA exposure is associated with a progressive increase in the number of smaller LDs within organoids. Sample sizes: vehicle *n* = 4, OA 200 μM *n* = 9, OA 400 μM *n* = 11, LA 200 μM *n* = 3, LA 400 μM *n* = 3.

These qualitative differences were reflected in quantitative LD-level metrics [Figs. 8(c), (d)]. In OA (400 µM)-treated organoids, mean LD volume increased during the early phase of exposure and continued to rise over time, consistent with a shift toward volume-dominated accumulation [Fig. 8(c)]. In contrast, LA (400 µM)-treated organoids exhibited a sustained increase in LD count throughout the 24-h observation window, whereas mean LD volume increased only modestly at early time points and subsequently plateaued or declined (see Supplementary Fig. S3 for the corresponding unnormalized values). At the lower dose (200 µM), OA- and LA-treated organoids did not show the same pronounced divergence between LD count and mean LD volume.

Mean LD dry mass followed similar temporal patterns to mean LD volume within each condition, consistent with OA and LA differing primarily in whether lipid burden is accommodated by droplet enlargement versus formation of additional droplets. To further assess LD biophysical properties, we quantified LD concentration and sphericity across all conditions [Fig. 8(d)]. LD concentration showed modest condition-dependent differences, with LA-treated LDs tending to exhibit slightly higher values than OA-treated LDs. LD sphericity remained near unity across conditions, indicating largely preserved overall droplet shape during the 24-h time window. LD size heterogeneity was further quantified using the coefficient of variation (CV) of individual LD volumes [Fig. 8(e)]. OA treatment led to a progressive increase in CV over time, indicating growing heterogeneity in LD size distribution. In contrast, LA-treated organoids showed relatively stable CV values, reflecting a comparatively more uniform accumulation across many droplets.

Together, these analyses reveal two distinct temporal modes of LD accumulation in response to unsaturated fatty acids. OA-treated organoids primarily increased total LD content through preferential enlargement of a smaller number of LDs, resulting in greater size heterogeneity, whereas LA-treated organoids accumulated lipids through sustained increases in LD number, yielding a more uniform size distribution of smaller droplets. These contrasting accumulation modes, summarized schematically in Fig. 8(f), provide a conceptual framework for interpreting fatty acid–dependent lipid storage strategies. To further examine whether the distinct LD accumulation modes observed in Fig. 8 are associated with differences in the 3D spatial distribution of LDs, we performed an exploratory centroid-based spatiotemporal analysis (Fig. S4 in the Supplementary Material). Specifically, we quantified LD spatial organization using three complementary centroid-based metrics that capture heterogeneity across local, global-network, and mesoscopic length scales (Gap NND CV, MST edge CV, and Grid Count IOD; see Fig. S4 for definitions). Collectively, these metrics suggest that the pronounced increase in LD count under LA treatment coincided with increasing spatial heterogeneity over time, whereas these spatial metrics change more modestly in OA-treated organoids.

## 4. Discussion and Conclusions

In this study, we established a quantitative, label-free framework for tracking LD dynamics in living mHOs using HT. Unlike endpoint-based or fluorescence-dependent assays, this approach enables continuous 3D observation of LD formation, growth, and redistribution under native conditions without perturbing lipid metabolism. By integrating HT with a depth-adaptive segmentation and spatiotemporal analysis pipeline, we quantitatively resolved LD accumulation across multiple scales—from individual droplets to intact organoids—under different FFA environments.

This work establishes a generalizable approach for non-invasive, longitudinal quantification of LD dynamics in 3D organoid systems. By linking droplet-level behavior to organoid-scale kinetics and morphology, our framework provides insight into how fatty-acid class shapes lipid storage strategies and offers a tool for studying metabolic remodeling in physiologically relevant models.

Population-level analyses demonstrated that both OA and LA induce robust, concentration-dependent LD accumulation in mHOs while largely preserving organoid integrity (Fig. 7). However, single–LD–resolved analyses revealed that similar increases in total LD burden arise through fundamentally different temporal strategies depending on fatty-acid class (Fig. 8). OA-treated organoids primarily accumulated lipids through preferential enlargement of a smaller number of LDs. This mode was characterized by early increases in LD volume accompanied by transient stabilization or reduction in LD number at later time points, leading to increased size heterogeneity. Such volume-dominated accumulation is consistent with extensive evidence that OA potently induces LD hypertrophy and enlargement in diverse cell types^55–57^. In contrast, LA-treated organoids accumulated lipids predominantly through sustained increases in LD number, yielding a more uniform population of smaller droplets. This number-dominated accumulation may be consistent with the biochemical properties of polyunsaturated fatty acids (PUFAs), which are more susceptible to lipid peroxidation and are thought to be preferentially sequestered into neutral lipid stores as a protective mechanism^58,59^.

Importantly, these divergent accumulation modes were not readily apparent from organoid-level metrics alone, underscoring the value of resolving LD dynamics at the single-droplet level. Together, our results demonstrate that fatty-acid class dictates not only the magnitude of lipid storage but also the temporal logic by which LDs are generated and remodeled in 3D organoids.

Technically, this work addresses a key limitation of 3D RI-based LD analysis. In thick organoids, depth-dependent RI variations and optical aberrations render single-threshold segmentation unreliable. Our adaptive multi-threshold segmentation strategy mitigates these effects by incorporating local contrast and morphological constraints, enabling robust LD detection across depth layers. This capability is critical for accurate longitudinal analysis of LD remodeling in intact organoids.

Nevertheless, several limitations remain. First, RI-based volume estimates may be influenced by axial elongation due to anisotropic resolution and aberrations. As with all limited-angle HT implementations, axial resolution is reduced relative to lateral resolution due to the missing-cone problem, which can bias absolute measurements of object shape and sphericity. Second, missing-cone artifacts can bias absolute RI values, affecting derived dry-mass and concentration estimates. Third, segmentation accuracy remains dependent on image quality, particularly in deeper regions where high-RI non-lipid structures may confound LD identification. Future improvements could include adaptive RI correction, confidence-weighted segmentation, and calibration strategies to further enhance quantitative accuracy^60–62^.

Beyond methodological advances, this framework provides new opportunities for studying lipid metabolism and disease-relevant phenotypes in organoids. Clinically, LD size, number, and spatial distribution are key determinants distinguishing microvesicular and macrovesicular steatosis, which are associated with different liver pathologies^2,63,64^. The ability to quantify LD dynamics in three dimensions over time enables discrimination between stable lipid sequestration and dynamic lipid turnover—an important distinction in metabolic disorders such as steatosis, obesity, diabetes, and cancer^65–67^. Moreover, the non-invasive and label-free nature of this approach makes it well suited for longitudinal drug screening and metabolic perturbation studies, where repeated measurements of the same organoid are essential^68–70^. Although the present study focused on hepatic organoids, extension of this framework to hepatocyte-enriched or vascularized multi-lineage organoids may provide deeper insight into LD remodeling in more physiologically relevant contexts^71,72^.

In summary, our results demonstrate that LD accumulation in organoids is a dynamic, multi-scale remodeling process shaped by fatty-acid class. By combining HT with adaptive segmentation and single-droplet analysis, we provide a generalizable framework for quantitatively linking LD biogenesis and morphology in living 3D systems.

## Disclosures

H.L., and Y.K.P. have financial interests in Tomocube, a company that commercializes HT instruments.

## Code and Data Availability

Representative HT 3D time-lapse datasets used in this study are publicly available. To ensure transparency and reproducibility while keeping data volume manageable, we provide one representative dataset for each experimental group (control, OA-treated, and LA-treated organoids), including raw reconstructed RI tomograms and associated metadata required to reproduce all quantitative analyses shown in the main figures. All datasets are available at: https://zenodo.org/records/18251200.

All core analysis code used for LD segmentation, depth-adaptive thresholding, single-droplet tracking, and quantitative metric extraction is openly available at: https://github.com/BMOLKAIST/Organoid_LD. The repository includes detailed documentation and example scripts that reproduce the analyses performed on the representative datasets. Custom scripts used solely for figure formatting are described in the repository but are not required to reproduce the reported quantitative results.

## Acknowledgements

This work was supported by National Research Foundation of Korea grant funded by the Korea government (MSIT) (RS-2024-00442348, RS-2022-NR068141), Korea Institute for Advancement of Technology (KIAT) through the International Cooperative R&D program (P0028463), and Commercialization Promotion Agency for R&D Outcomes (COMPA) funded by the Ministry of Science and ICT(MSIT) (RS-2024-00440577).

## References

1. J. A. Olzmann, P. Carvalho, “Dynamics and functions of lipid droplets,” Nat. Rev. Mol. Cell Biol. 20(3), 137– 155 (2019) [doi:10.1038/s41580-018-0085-z].

2. D. Marti-Aguado et al., “Digital pathology and lipid droplet size as a key determinant of discrepancies between histology and MRI gradings in steatotic liver disease,” Eur. Radiol. (2025) [doi:10.1007/s00330-025-11919-0].

3. W.-Q. Leow et al., “Non-alcoholic fatty liver disease: the pathologist’s perspective,” Clin. Mol. Hepatol. 29(Suppl), S302–S318 (2023) [doi:10.3350/cmh.2022.0329].

4. D. Hendriks et al., “Engineered human hepatocyte organoids enable CRISPR-based target discovery and drug screening for steatosis,” Nat. Biotechnol. 41(11), 1567–1581 (2023) [doi:10.1038/s41587-023-01680-4].

5. E. Yildiz et al., “Hepatic lipid overload triggers biliary epithelial cell activation via E2Fs,” eLife 12, e81926 (2023) [doi:10.7554/eLife.81926].

6. M. N. B. Ramli et al., “Human Pluripotent Stem Cell-Derived Organoids as Models of Liver Disease,” Gastroenterology 159(4), 1471-1486.e12 (2020) [doi:10.1053/j.gastro.2020.06.010].

7. M. Kozyra et al., “Human hepatic 3D spheroids as a model for steatosis and insulin resistance,” Sci. Rep. 8(1), 14297 (2018) [doi:10.1038/s41598-018-32722-6].

8. Y. Guan et al., “Human hepatic organoids for the analysis of human genetic diseases,” JCI Insight 2(17), (2023) [doi:10.1172/jci.insight.94954].

9. H. Kim et al., “Recent advances in label-free imaging and quantification techniques for the study of lipid droplets in cells,” Curr. Opin. Cell Biol. 87, 102342 (2024) [doi:10.1016/j.ceb.2024.102342].

10. R. Keshara, Y. H. Kim, A. Grapin-Botton, “Organoid Imaging: Seeing Development and Function,” Annu. Rev. Cell Dev. Biol. 38(1), 447–466 (2022) [doi:10.1146/annurev-cellbio-120320-035146].

11. H. Khadem et al., “Polarization-Sensitive Holotomography for Multidimensional Label-Free Imaging and Characterization of Lipid Droplets in Cancer Cells,” Adv. Sci. 12(42), e09420 (2025) [doi:10.1002/advs.202509420].

12. P. P. Laissue et al., “Assessing phototoxicity in live fluorescence imaging,” Nat. Methods 14(7), 657–661 (2017) [doi:10.1038/nmeth.4344].

13. Y. Liu et al., “Engineered Hydrogels for Organoid Models of Human Nonalcoholic Fatty Liver Disease,” Adv. Sci. 12(22), e17332 (2025) [doi:10.1002/advs.202417332].

14. G. Kim et al., “Holotomography,” Nat. Rev. Methods Primer 4(1), 51 (2024) [doi:10.1038/s43586-024-00327-1].

15. K. Kim et al., “Three-dimensional label-free imaging and quantification of lipid droplets in live hepatocytes,” Sci. Rep. 6(1), 36815 (2016) [doi:10.1038/srep36815].

16. D. Park et al., “Cryobiopsy: A Breakthrough Strategy for Clinical Utilization of Lung Cancer Organoids,” Cells 12(14), 1854 (2023) [doi:10.3390/cells12141854].

17. K. Han et al., “Gelatin-Based Soft-Tissue Sarcoma Organoids Recapitulate Patient Tumor Characteristics,” Biomater. Res. 29, 0293 (2025) [doi:10.34133/bmr.0293].

18. D. Kong et al., “In Vitro Modeling of Atherosclerosis Using iPSC-Derived Blood Vessel Organoids,” Adv. Healthc. Mater. 14(1), 2400919 (2025).

19. H. Kim et al., “Diverse bat organoids provide pathophysiological models for zoonotic viruses,” Science 388(6748), 756–762 (2025) [doi:10.1126/science.adt1438].

20. J. Rosen et al., “Roadmap on computational methods in optical imaging and holography [invited],” Appl. Phys. B 130(9), 166 (2024) [doi:10.1007/s00340-024-08280-3].

21. Y. Liu, S. Uttam, “Perspective on quantitative phase imaging to improve precision cancer medicine,” J. Biomed. Opt. 29(S2), S22705 (2024) [doi:10.1117/1.JBO.29.S2.S22705].

22. D. Pirone et al., “Stain-free identification of cell nuclei using tomographic phase microscopy in flow cytometry,” Nat. Photonics 16(12), 851–859 (2022) [doi:10.1038/s41566-022-01096-7].

23. D. Jin et al., “Tomographic phase microscopy: principles and applications in bioimaging,” J. Opt. Soc. Am. B 34(5), B64–B77 (2017).

24. A. Marzi et al., “Quantitative Phase Imaging as Sensitive Screening Method for Nanoparticle-Induced Cytotoxicity Assessment,” Cells 13(8), 697 (2024) [doi:10.3390/cells13080697].

25. P. Anantha et al., “Uncovering Astrocyte Morphological Dynamics Using Optical Diffraction Tomography and Shape-Based Trajectory Inference,” Adv. Healthc. Mater. 14(6), 2402960 (2025) [doi:10.1002/adhm.202402960].

26. H. Hugonnet, M. Lee, Y. Park, “Optimizing illumination in three-dimensional deconvolution microscopy for accurate refractive index tomography,” Opt. Express 29(5), 6293–6301 (2021) [doi:10.1364/OE.412510].

27. Y. Chung et al., “Fourier space aberration correction for high resolution refractive index imaging using incoherent light,” Opt. Express 32(11), 18790–18799 (2024) [doi:10.1364/OE.518479].

28. H. Hugonnet, M. J. Lee, Y. K. Park, “Quantitative phase and refractive index imaging of 3D objects via optical transfer function reshaping,” Opt. Express 30(8), 13802–13809 (2022) [doi:10.1364/OE.454533].

29. M. J. Lee et al., “Long-term three-dimensional high-resolution imaging of live unlabeled small intestinal organoids via low-coherence holotomography,” Exp. Mol. Med. 56(10), 2162–2170 (2024) [doi:10.1038/s12276-024-01312-0].

30. J. Cho et al., “Label-free, High-Resolution 3D Imaging and Machine Learning Analysis of Intestinal Organoids via Low-Coherence Holotomography,” J. Vis. Exp. JoVE(222), e68529 (2025) [doi:10.3791/68529].

31. E. Y. Jeong et al., “Label-free long-term measurements of adipocyte differentiation from patient-driven fibroblasts and quantitative analyses of in situ lipid droplet generation,” JOSA A 41(11), C125–C136 (2024) [doi:10.1364/JOSAA.528703].

32. S. Park et al., “Label-Free Tomographic Imaging of Lipid Droplets in Foam Cells for Machine-Learning-Assisted Therapeutic Evaluation of Targeted Nanodrugs,” ACS Nano 14(2), 1856–1865 (2020) [doi:10.1021/acsnano.9b07993].

33. C. Oh et al., Morphology-Preserving Holotomography: Quantitative Analysis of 3D Organoid Dynamics, arXiv (2025) [doi:10.48550/arXiv.2512.22486].

34. M. Ricchi et al., “Differential effect of oleic and palmitic acid on lipid accumulation and apoptosis in cultured hepatocytes,” J. Gastroenterol. Hepatol. 24(5), 830–840 (2009) [doi:10.1111/j.1440-1746.2008.05733.x].

35. A. Eynaudi et al., “Differential Effects of Oleic and Palmitic Acids on Lipid Droplet-Mitochondria Interaction in the Hepatic Cell Line HepG2,” Front. Nutr. 8, (2021) [doi:10.3389/fnut.2021.775382].

36. F. S. Teixeira et al., “Differential Lipid Accumulation on HepG2 Cells Triggered by Palmitic and Linoleic Fatty Acids Exposure,” Molecules 28(5), 2367 (2023) [doi:10.3390/molecules28052367].

37. M. Zhang et al., “The Different Mechanisms of Lipid Accumulation in Hepatocytes Induced by Oleic Acid/Palmitic Acid and High-Fat Diet,” Molecules 28(18), (2023) [doi:10.3390/molecules28186714].

38. R. Barer, “Determination of Dry Mass, Thickness, Solid and Water Concentration in Living Cells,” Nature 172(4389), 1097–1098 (1953) [doi:10.1038/1721097a0].

39. H. Kim et al., “Recent advances in label-free imaging and quantification techniques for the study of lipid droplets in cells,” Curr. Opin. Cell Biol. 87, 102342 (2024) [doi:10.1016/j.ceb.2024.102342].

40. G. Popescu et al., “Optical imaging of cell mass and growth dynamics,” Am. J. Physiol.-Cell Physiol. 295(2), C538–C544 (2008) [doi:10.1152/ajpcell.00121.2008].

41. N. A. Ducharme, P. E. Bickel, “Minireview: Lipid Droplets in Lipogenesis and Lipolysis,” Endocrinology 149(3), 942–949 (2008) [doi:10.1210/en.2007-1713].

42. G. Onal et al., “Lipid Droplets in Health and Disease,” Lipids Health Dis. 16(1), 128 (2017) [doi:10.1186/s12944-017-0521-7].

43. R. W. Klemm, P. Carvalho, “Lipid Droplets Big and Small: Basic Mechanisms That Make Them All,” Annu. Rev. Cell Dev. Biol. 40(Volume 40, 2024), 143–168 (2024) [doi:10.1146/annurev-cellbio-012624-031419].

44. A. J. Mathiowetz, J. A. Olzmann, “Lipid droplets and cellular lipid flux,” Nat. Cell Biol. 26(3), 331–345 (2024) [doi:10.1038/s41556-024-01364-4].

45. M. B. Schott et al., “Lipid droplet size directs lipolysis and lipophagy catabolism in hepatocytes,” J. Cell Biol. 218(10), 3320–3335 (2019) [doi:10.1083/jcb.201803153].

46. A. Kassan et al., “Acyl-CoA synthetase 3 promotes lipid droplet biogenesis in ER microdomains,” J. Cell Biol. 203(6), 985–1001 (2013) [doi:10.1083/jcb.201305142].

47. H. B. Castillo et al., “Oleic acid differentially affects lipid droplet storage of de novo synthesized lipids in hepatocytes and adipocytes,” Chem. Commun. 60(23), 3138–3141 (2024) [doi:10.1039/D3CC04829B].

48. T. Plötz et al., “The role of lipid droplet formation in the protection of unsaturated fatty acids against palmitic acid induced lipotoxicity to rat insulin-producing cells,” Nutr. Metab. 13(1), 16 (2016) [doi:10.1186/s12986-016-0076-z].

49. S. Schuster et al., “Oxidized linoleic acid metabolites induce liver mitochondrial dysfunction, apoptosis, and NLRP3 activation in mice,” J. Lipid Res. 59(9), 1597–1609 (2018) [doi:10.1194/jlr.M083741].

50. L.-J. Su et al., “Reactive Oxygen Species-Induced Lipid Peroxidation in Apoptosis, Autophagy, and Ferroptosis,” Oxid. Med. Cell. Longev. 2019, 5080843 (2019) [doi:10.1155/2019/5080843].

51. Y. Zheng et al., “Emerging mechanisms of lipid peroxidation in regulated cell death and its physiological implications,” Cell Death Dis. 15(11), 859 (2024) [doi:10.1038/s41419-024-07244-x].

52. A. Hess et al., “Single-cell transcriptomics stratifies organoid models of metabolic dysfunction-associated steatotic liver disease,” EMBO J. 42(24), e113898 (2023) [doi:10.15252/embj.2023113898].

53. U. Ziegler, P. Groscurth, “Morphological Features of Cell Death,” Physiology 19(3), 124–128 (2004) [doi:10.1152/nips.01519.2004].

54. S. Elmore, “Apoptosis: A Review of Programmed Cell Death,” Toxicol. Pathol. 35(4), 495–516 (2007) [doi:10.1080/01926230701320337].

55. S. Murphy, S. Martin, R. G. Parton, “Quantitative Analysis of Lipid Droplet Fusion: Inefficient Steady State Fusion but Rapid Stimulation by Chemical Fusogens,” PLOS ONE 5(12), e15030 (2010) [doi:10.1371/journal.pone.0015030].

56. Y. Zheng et al., “Mitochondrial metabolic dysfunction and non-alcoholic fatty liver disease: new insights from pathogenic mechanisms to clinically targeted therapy,” J. Transl. Med. 21(1), 510 (2023) [doi:10.1186/s12967-023-04367-1].

57. L. L. Listenberger et al., “Triglyceride accumulation protects against fatty acid-induced lipotoxicity,” Proc. Natl. Acad. Sci. 100(6), 3077–3082 (2003) [doi:10.1073/pnas.0630588100].

58. T. Petan, E. Jarc, M. Jusović, “Lipid Droplets in Cancer: Guardians of Fat in a Stressful World,” Molecules 23(8), 1941 (2018) [doi:10.3390/molecules23081941].

59. A. P. Bailey et al., “Antioxidant Role for Lipid Droplets in a Stem Cell Niche of Drosophila,” Cell 163(2), 340– 353 (2015) [doi:10.1016/j.cell.2015.09.020].

60. H. Hugonnet et al., “Pupil phase series: a fast, accurate, and energy-conserving model for forward and inverse light scattering in thick biological samples,” Opt. Express 33(16), 34255–34266 (2025) [doi:10.1364/OE.568019].

61. C. Oh et al., “Digital aberration correction for enhanced thick tissue imaging exploiting aberration matrix and tilt-tilt correlation from the optical memory effect,” Nat. Commun. 16(1), 1685 (2025) [doi:10.1038/s41467-025-56865-z].

62. K.-S. Chuang et al., “Fuzzy c-means clustering with spatial information for image segmentation,” Comput. Med. Imaging Graph. 30(1), 9–15 (2006) [doi:10.1016/j.compmedimag.2005.10.001].

63. M. N. B. Kristiansen et al., “Molecular Characterization of Microvesicular and Macrovesicular Steatosis Shows Widespread Differences in Metabolic Pathways,” Lipids 54(1), 109–115 (2019) [doi:10.1002/lipd.12121].

64. E. M. Brunt, “Pathology of fatty liver disease,” Mod. Pathol. 20, S40–S48 (2007) [doi:10.1038/modpathol.3800680].

65. J. Taylor et al., “Generation of immune cell containing adipose organoids for in vitro analysis of immune metabolism,” Sci. Rep. 10(1), 21104 (2020) [doi:10.1038/s41598-020-78015-9].

66. X. Li et al., “Research Progress on Lipophagy-Mediated Exercise Intervention in Non-Alcoholic Fatty Liver Disease,” Int. J. Mol. Sci. 25(6), 3153 (2024) [doi:10.3390/ijms25063153].

67. D. E. Berardi et al., “Lipid droplet turnover at the lysosome inhibits growth of hepatocellular carcinoma in a BNIP3-dependent manner,” Sci. Adv. 8(41), eabo2510 (2022) [doi:10.1126/sciadv.abo2510].

68. P. J. Tebon et al., “Drug screening at single-organoid resolution via bioprinting and interferometry,” Nat. Commun. 14(1), 3168 (2023) [doi:10.1038/s41467-023-38832-8].

69. C. E. Serafini et al., “Non-invasive label-free imaging analysis pipeline for in situ characterization of 3D brain organoids,” Sci. Rep. 14(1), 22331 (2024) [doi:10.1038/s41598-024-72038-2].

70. I. Abd El-Sadek et al., “Label-free visualization and quantification of the drug-type-dependent response of tumor spheroids by dynamic optical coherence tomography,” Sci. Rep. 14(1), 3366 (2024) [doi:10.1038/s41598-024-53171-4].

71. T. Takebe et al., “Vascularized and Complex Organ Buds from Diverse Tissues via Mesenchymal Cell-Driven Condensation,” Cell Stem Cell 16(5), 556–565 (2015) [doi:10.1016/j.stem.2015.03.004].

72. T. Takebe et al., “Vascularized and functional human liver from an iPSC-derived organ bud transplant,” Nature 499(7459), 481–484 (2013) [doi:10.1038/nature12271].

